# The potential impacts of vector host species fidelity on zoonotic arbovirus transmission

**DOI:** 10.1101/2024.05.08.593124

**Authors:** Tijani A. Sulaimon, Anthony J. Wood, Michael B. Bonsall, Michael Boots, Jennifer S. Lord

## Abstract

The interaction between vector host preference and host availability on vector blood feeding behaviour has important implications for the transmission of vector-borne pathogens. In particular in multi-host disease systems the fidelity of the vector biting behaviour has the potential to have important implications to disease outcomes, particularly when there are amplifying and dead-end hosts. Using a mathematical model we showed that vector fidelity to the host species they take a first blood meal from leads to non-homogeneous mixing between hosts and vectors. Taking Japanese encephalitis virus (JEV) as a case study, we investigated how vector preference for amplifying vs dead-end hosts and fidelity can influence JEV transmission. We show that in regions where pigs (amplifying hosts) are scarce compared to cattle (dead-end hosts preferred by common JEV vectors), JEV can still be maintained through vector fidelity. Our findings demonstrate the importance of considering fidelity as a potential driver of transmission, particularly in scenarios such as Bangladesh and India where the composition of the host community might initially suggest that transmission is not possible.

## INTRODUCTION

It is estimated that there are *>* 14 000 species of hematophagous arthropods (Ribeiro, 1995), some of which are vectors of vertebrate pathogens. Arthropod vectors have evolved, to a greater or lesser degree, particular preferences for certain host species, or groups of hosts, from which to take a blood meal (Besansky et al., 2004; Takken and Verhulst, 2013). Here we define host preference as the genetically based tendency to respond to particular host cues (Clements et al., 1999). During host-seeking, genetically determined host preference combines primarily with relative host availability and vector density, to determine realised host choice. For example, Gurtler et al. (1997) showed that triatomines fed less frequently on humans when dogs or chickens were present in bedroom areas and when triatomine density was higher. Similarly, Thiemann et al. (2011) demonstrated that temporal changes in the blood feeding patterns of the mosquito *Culex tarsalis*, from ardeid birds in the summer to mammals once ardeids had fledged and left the area, were driven by changes in host availability and mosquito abundance in addition to underlying preference.

The interaction between vector host preference and host availability on vector blood feeding behaviour has consequences for the transmission dynamics of vector-borne pathogens. Miller and Huppert (2013), using a general modelling framework, showed that higher host diversity can result in amplification of a pathogen when the preferred host is also the host with highest competence. However, diversity can cause dilution when a vector prefers the host that is of lower competence, or when a vector does not have an underlying preference for any specific host species. Researchers have also carried out analyses for specific pathogens and contexts with respect to the effect of host choice. Killeen et al. (2001) used models parameterised by field data to show that increased cattle populations could reduce malaria transmission in The Gambia but not in Tanzania due to higher *Anopheles* preference for cattle compared with humans in The Gambia. Ginsberg et al. (2021) showed that latitudinal differences in tick host-seeking behaviour contributes to the geographic pattern of Lyme disease in the eastern United States. Similarly, Hamer et al. (2011) showed that spatial variation in West Nile virus transmission can in part be explained by variation in vector host choice.

In most analyses, it is assumed that the relationship between vector host choice and relative host availability is linear. Different functional responses of vectors to relative host availability have, however, been modelled by Yakob and Walker (2016). Yakob and Walker (2016) demonstrated that the assumed function can have implications for predicted effects of control measures aimed at reducing vector biting rates. Models have also been used to show that vector preference for infected hosts relative to susceptible hosts can have important effects on disease dynamics King-solver (1987). Despite this range of modelling efforts, to our knowledge, there have been few analyses of the possible effects of vector memory and host fidelity on host choice and, therefore, pathogen transmission dynamics. We are only aware of two modelling studies that consider the potential for mosquitoes to alter their host choice based on previous feeding experience (Sardar et al., 2015a,b).

Sardar et al. (2015b) proposed a model of dengue virus transmission between human and mosquito populations using “fractional-order” (FO) differential equations, which incorporate memory through the order of the fractional derivative. In systems governed by integer-order differential equations, the future evolution depends only on its present state (i.e. Markovian). The future state of FO models, on the other hand, can depend on both the present and past states, with the order of the fractional derivative being the memory index. In their model, they incorporated a memory index into the mosquito biting rate, which reduces the contact rate between hosts and mosquitoes by increasing human awareness of dengue transmission and influencing mosquito behaviour towards host choice and location. Their results suggest that as the capacity of a system to retain memory increases, the probability of dengue persistence in a population will also increase. Sardar et al. (2015a) compared three models of dengue transmission: one based on ordinary differential equations and two based on FO differential equations, with both FO models considering memory in mosquito biting rate and population abundance. The first FO model assumed similar FO dynamics for mosquitoes and humans (equal memory indices), while the other FO model assumed they were different. The proposed models were validated using published monthly dengue incidence data from two provinces of Venezuela during the period 1999–2002. The results indicate that the memory and learning behaviour of mosquitoes will increase the transmission of mosquito-borne diseases and have a considerable impact on mosquito recruitment rate. Furthermore, the model with memory in both the host and vector populations is more consistent with dengue epidemic data. Both of these modelling studies assumed a single host species (humans) and, therefore, did not explicitly model vector preference for specific host species and the effect of fidelity behaviour on host choice. Also, it is not clear how to measure the memory index experimentally.

There is now substantial empirical evidence that vectors can learn from previous experience (Vinauger et al., 2016). For mosquitoes, this is both with respect to choice of oviposition sites and hosts. There is evidence that both attraction and aversion behaviour can be learned when mosquitoes are exposed to a particular odour alongside either a positive or negative experience (Vinauger et al., 2014, 2018).

We are not aware of any modelling studies that consider the potential effects of vector fidelity on the transmission dynamics of zoonotic vector-borne pathogens. Given that, by necessity, the community of vector species that transmit zoonotic pathogens must feed on multiple host species, the question of the effect of learning during host choice on their transmission dynamics is particularly pertinent.

Our main goal was to fill this research gap by introducing a model where vectors become “imprinted” to a host species on their first blood meal and have a degree of fidelity to them for future feeds. This extends the traditional compartmental model (Brauer, 2008; McDonald, 1957) to have a “memory” parameter. We use this to assess the impact of fidelity on transmission dynamics between vector and host using analytic and numerical methods.

We use Japanese encephalitis virus (JEV) as a case study for this model. Japanese encephalitis is a zoonotic disease caused by (JEV Solomon et al. (2000)), primarily transmitted by mosquitoes (Gresser et al., 1958; Mackenzie et al., 2004). It poses a significant health threat, particularly in rural areas of Asia where rice cultivation and pig farming are prevalent (Misra and Kalita, 2010). There is no specific treatment available for JE, but the human disease is preventable by vaccination (Turtle and Solomon, 2018).

Futhermore, the transmission of JEV involves a cycle that occurs between multiple species of mosquitoes and reservoir hosts, including pigs and birds (Bukscher et al., 1959; Gresser et al., 1958; Misra and Kalita, 2010). To maintain the enzootic cycle of the JEV, a mosquito must become infected by biting an infectious host and subsequently survive, develop a sufficient viral load in the saliva for transmission, find, and bite a susceptible host. For an infected host to transmit the virus to a mosquito vector, it must attain an adequate concentration of the virus in its bloodstream.

Most importantly, two studies have provided evidence for host fidelity in potential vectors of JEV. Mwandawiro et al. (1999, 2000) showed through field experiments that although *Culex vishnui, Culex tritaeniorhynchus* and *Culex gelidus* would more frequently choose a cow over a pig in host choice experiments, they had a tendency to take a subsequent blood meal on the host species from which they had previously fed. Those initially attracted to pigs were more likely to take subsequent blood meals from pigs. As such, JE is an excellent system in which to examine the impact of vector fidelity on infectious disease outcomes.

Intensive pig farming is associated with the prevalence of JE in regions such as China and Japan (Lord et al., 2015; Misra and Kalita, 2010). Pigs are the main hosts for JEV amplification and may re-transmit to susceptible feeding mosquitoes, whereas cattle are considered dead-end hosts and do not contribute to JEV transmission. Hence, high relative availability of cattle compared with pigs may dilute JEV transmission due to the preference of JEV vectors for this host. However, JE cases can still occur in regions with low pig density compared to cattle, such as certain areas of Bangladesh and India (Lord et al., 2015). For example, in three JE-endemic districts of Rajshahi Division, Bangladesh, the ratio of cattle to pigs is approximately 140:1. Similarly, in some JE-endemic regions of India, cattle can outnumber pigs by a ratio of up to 20:1. In these regions where there is a relatively high density of cattle, which are preferred by *Culex tritaeniorhynchus* mosquitoes, could the fidelity behaviour of mosquitoes explain the persistence of JEV?

## MATERIALS AND METHODS

### A vector-borne pathogen model with host preference and fidelity

We describe a model of pathogen transmission that involves two host species and a vector. This is a general compartmental model, though in this work we consider model parameters so as to capture the dynamics of JEV over host populations of pigs and cows and a mosquito vector population.

### Fidelity-free transmission dynamics

To begin, the host population comprises two distinct groups; *amplifying* hosts (*A*) and *dead-end* hosts (*D*). The amplifying host population (*H*_*A*_) is capable of replicating and transmitting the virus to vectors. The dead-end host population (*H*_*D*_) can be infected but does not contribute to transmission. For each host species *i* ∈ {*A, D*}, we categorise into three compartments: susceptible (*S*_*i*_), infected (*I*_*i*_), and recovered (*R*_*i*_) individuals, with the total population given by *H*_*i*_ = *S*_*i*_ + *I*_*i*_ + *R*_*i*_. We assume the infection is nonfatal and all infected hosts recover at rate *γ*. For simplicity, the birth and death rates of both hosts are fixed at *µ*, preserving the total population in time.

The mosquito population follows a susceptible-infected dynamic (SI). Mosquitoes are introduced at a rate *µ*_*m*_ into the susceptible population *S*^*m*^. Mosquitoes die at rate *µ*_*m*_, preserving the total population, and bite hosts at a rate *α*. The mosquito’s choice of host type is determined (noting we are not yet considering fidelity) based on a host preference. We define the probability *ρ*_*i*_ that a mosquito chooses host species *i* for a particular blood meal as:

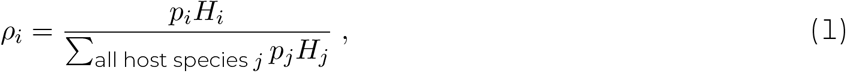

with *p*_*i*_ the genetically determined host preference for species *i* and *j* represents each individual host species.

If a susceptible mosquito bites an infected amplifying host (*I*_*A*_), that mosquito becomes infected with probability *β*. Similarly, if an infected mosquito (*I*^*m*^) bites a susceptible host (of any species), that host becomes infected with probability *β*. Susceptible mosquitoes, therefore, enter the infected compartment at rate 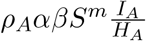. Hosts *i* enter the infected compartment at rate 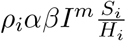.

Combining these infection dynamics, host recovery and the birth-death processes, the time evolution of this process can be described by the following set of ordinary differential equations:

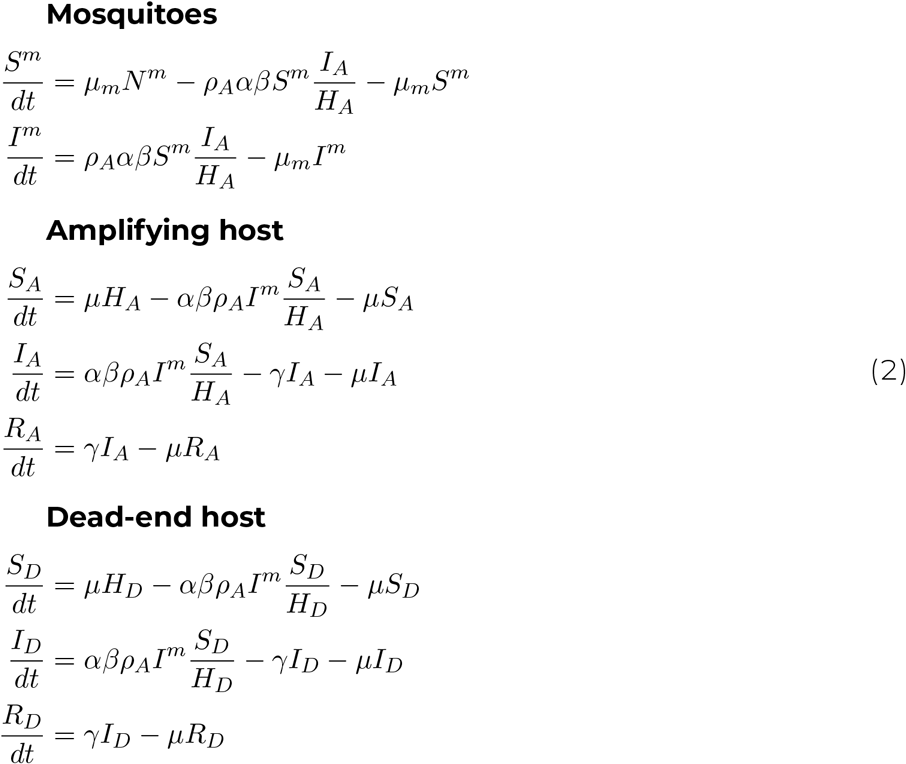

### Incorporating mosquito fidelity behaviour

The model described in Equation 2 assumes that the host choice of mosquitoes depends only on the relative abundance of each host, and the initial preference of mosquitoes (Equation 1). We now generalise this to include *fidelity*, where host choice is biased towards the host type that a mosquito *first* took a blood meal from (or is *imprinted* on).

We define *f* ∈ [0, 1] as the fidelity. For a mosquito imprinted on host *i*, the mosquito remains “loyal” to host *i* for any subsequent bite with probability *f* . If the mosquito does not remain loyal (probability 1 − *f*), then its choice of host is determined by the preference described in Equation 1. When *f* = 0 (no fidelity), mosquitoes behave solely based on host density and initial preferences. Conversely, for *f* = 1 (total fidelity), mosquitoes exclusively bite the host type that its first bite was taken from (Fig. 1). For subsequent bites by mosquitoes imprinted on host *i*, the proportion of bites on host *i* becomes *ρ*_*i*_ + *fρ*_*¬i*_, where *ρ*_*¬i*_ represents the proportion of first mosquito bites on the other host type. In contrast, for mosquitoes imprinted on the other host type, the proportion of subsequent bites on host *i* is (1 − *f*)*ρ*_*i*_.

**Figure 1:**
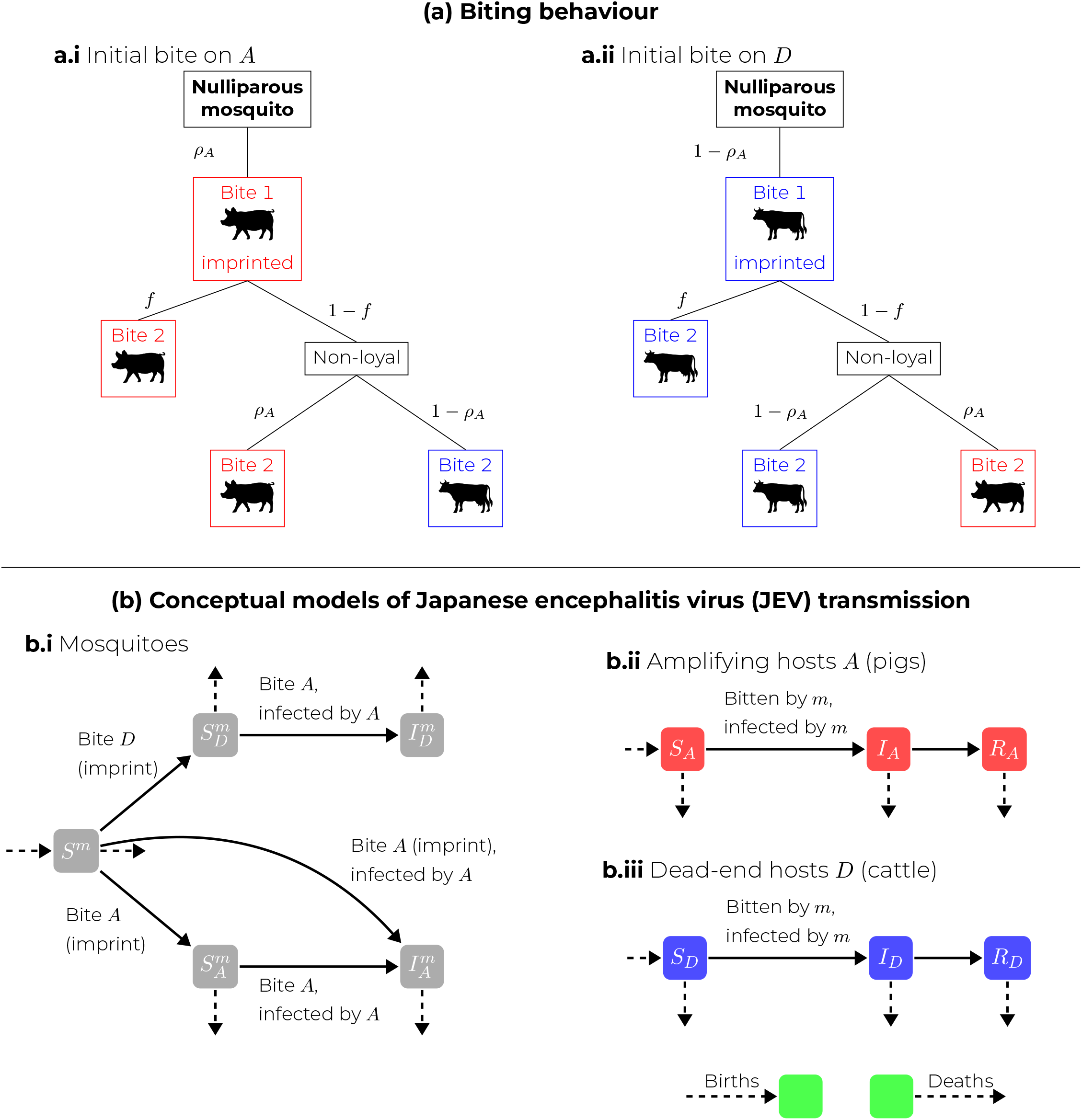
(a): Biting behaviour of mosquitoes, when (i) imprinted on an amplifying host *A*, and (ii) on a dead-end host *D*. A mosquito’s first bite is determined by host density and the host preference (modulated by the parameters *ρ*_*A*_, *ρ*_*D*_ = 1 − *ρ*_*A*_). All subsequent bites are determined by a mixture of fidelity (modulated by the parameter *f*), and host preference if the mosquito is not loyal to its imprinted host species. (b): Compartmental models. (i) Mosquitoes are introduced nulliparous (*S*^*m*^), and imprinted onto the host species they first bite 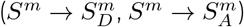, and later may become infected by biting infected amplifying hosts (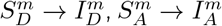, with potential for 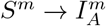 if infected on their imprinting bite). Mosquitoes can die in any compartment. (ii), (iii) Amplifying and dead-end hosts respectively. Hosts can become infected if bitten by an infected mosquito and later go on to recover. Hosts are born susceptible and can die in any compartment.

To generalise Equation 2 to include fidelity, we need to divide the population of mosquitoes into three subgroups: *nulliparous* (yet to take a blood meal) (*S*^*m*^), imprinted on an amplifying host 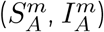, and imprinted on a dead-end host 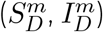. Susceptible mosquitoes imprinted on amplifying and dead-end hosts are recruited from the nulliparous group at rates 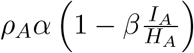 and *ρ*_*D*α_, respectively, where 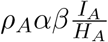 is the rate at which nulliparous mosquitoes become infected by taking their first blood meal on infected amplifying hosts. The rate at which susceptible mosquitoes imprinted on a host species *i* ∈ {*A, D*} become infected is 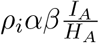. Incorporating this subclassification of mosquitoes, Equation 2 generalises to:

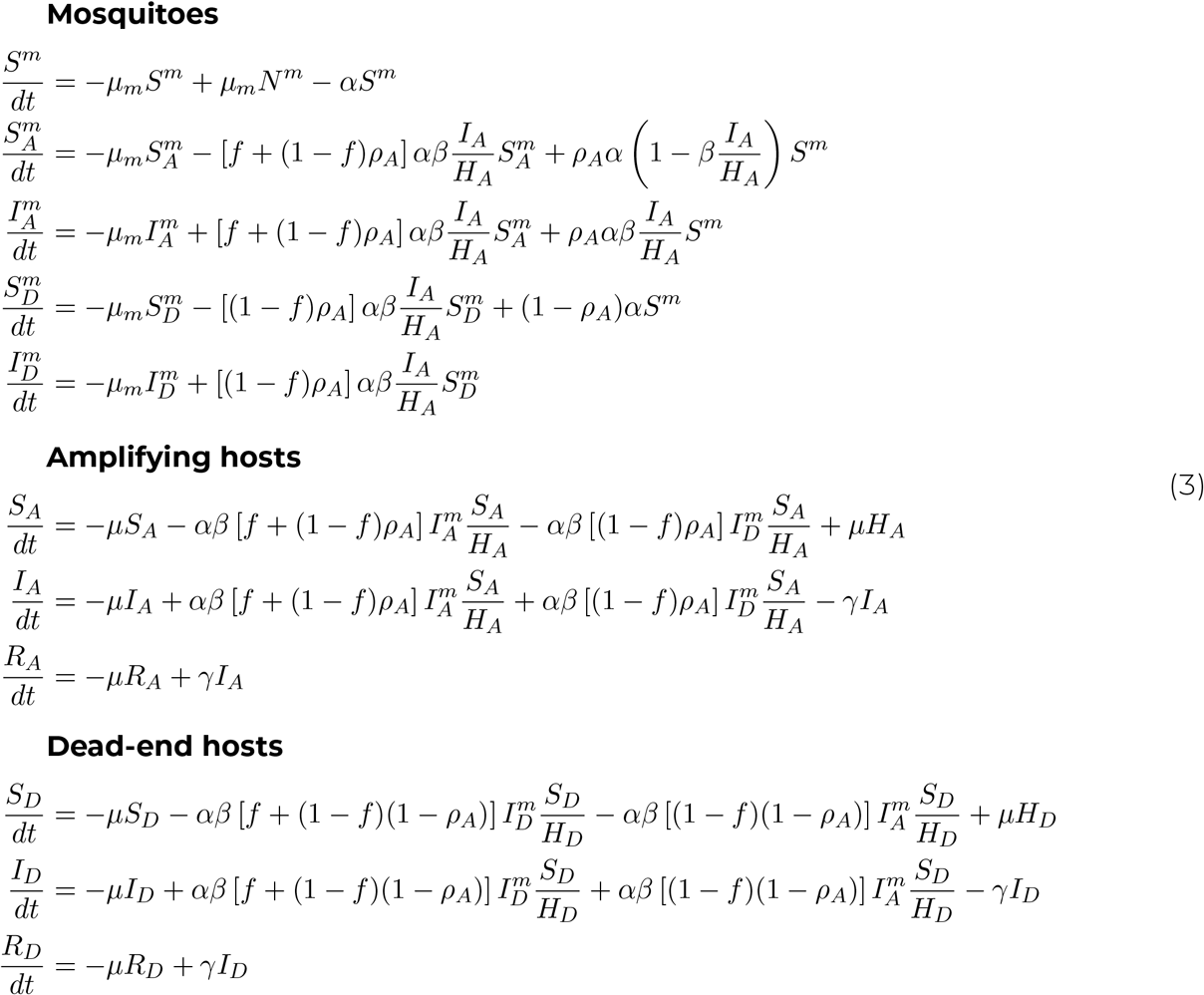

## RESULTS

### Host choice experiment

We first illustrate the implications of fidelity as described in Figure 1 on the host species choice of mosquitoes. Considering a host population comprising an *equal* proportion of cows (dead-end hosts, *D*) and pigs (amplifying hosts, *A*), Figure 2 shows the proportion of *initial* feeds on the two host species, and how the proportion of all *subsequent* feeds changes with fidelity. For no fidelity (*f* = 0, top row) where mosquitoes have no memory of the host they first fed on, the proportion of subsequent bites mirrors that of initial host preference. As the value for *f* increases, feeds are increasingly from mosquitoes imprinted onto that species, where with total fidelity (*f* = 1, bottom row) mosquitoes exclusively feed on the host to which they were imprinted. Note that the proportion of subsequent feeds per host remains constant.

**Figure 2:**
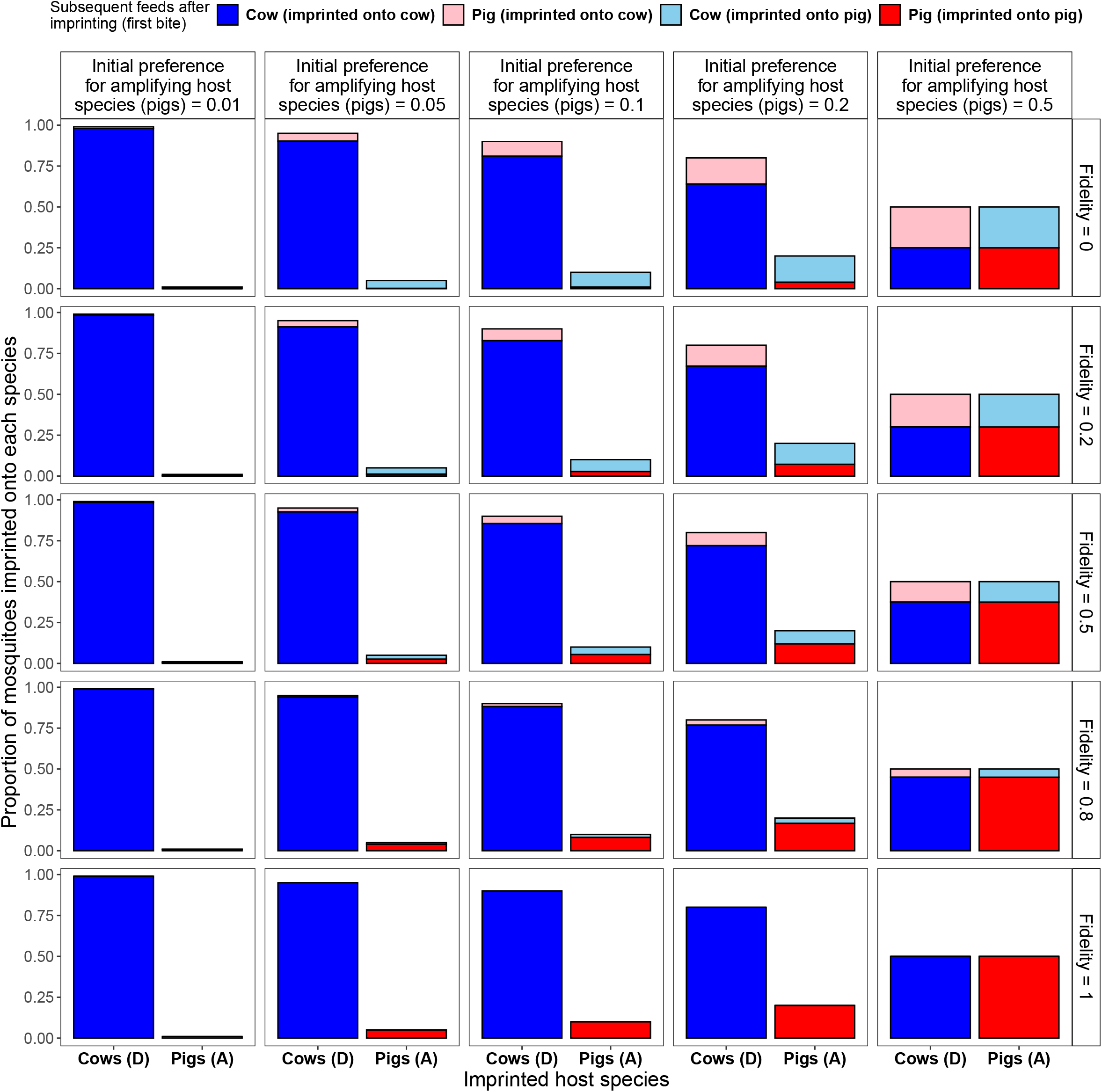
Variation of feeding patterns of mosquitoes with fidelity, in the instance where cows (D) and pigs (A) are available in equal proportions. The height of the bars indicates the proportion of mosquitoes that are imprinted on that host species (x-axis). The fill indicates the proportion of *subsequent bites* by those imprinted mosquitoes. With an increasing preference for pigs (left-to-right), a higher proportion of imprinting bites are on pigs. With increasing fidelity (top-to-bottom), the proportion of subsequent bites remaining loyal to the imprinted host increases. In the case of no fidelity (*f* = 0, top row), the proportion of subsequent bites replicates the proportion of initial bites. While the overall proportions feeding on D or A does not change with fidelity — as fidelity increases the population feeding on cattle vs pigs becomes increasingly isolated from each other.

### Basic reproduction number

From the full model in Equation 3, we derived an expression for the basic reproduction number (*R*_0_) in the case of an infection-free equilibrium, where populations are entirely susceptible. The basic reproduction number is a fundamental epidemiological metric that is used to assess the potential for pathogen transmission in a population. It represents the average number of secondary infections caused by a single infected individual in a fully susceptible population. Using the parameters described in Table 3, we analysed the influence of fidelity, host composition, and mosquito-to-host ratio on *R*_0_ (Figure 3).

**Table 1:** Definition of variables used in the model.

| Symbol | Meaning |
| --- | --- |
| $S_A$ | Number of susceptible individuals of amplifying host species |
| $I_A$ | Number of infected (infectious) individuals of amplifying host species |
| $S^m$ | Number of susceptible nulliparous mosquitoes |
| $S_A^m$ | Number of susceptible mosquitoes imprinted on amplifying host species |
| $I_A^m$ | Number of infectious mosquitoes imprinted on amplifying host species |
| $S_D^m$ | Number of susceptible mosquitoes imprinted on dead-end host species |
| $I_D^m$ | Number of infectious mosquitoes imprinted on dead-end host species |
| $N^m$ | Total mosquito population ($S^m + S_A^m + I_A^m + S_D^m + I_D^m$) (constant) |

**Table 2:** Definition of parameters used in the model.

| Symbol | Meaning |
| --- | --- |
| $H_A$ | Total number of individuals of amplifying host species (constant, $H_A = S_A + I_A + R_A$) [individual per area] |
| $H_D$ | Total number of individuals from dead-end host species (constant) [individual per area] |
| $\rho_i$ | Proportion of initial blood meals taken on species $i$ [unitless] |
| $\alpha$ | Individual mosquito biting rate [$\text{time}^{-1}$] |
| $\beta$ | Transmission probability: mosquito to host and host to mosquitoes [unitless] |
| $\gamma$ | Recovery rate for an individual host species (inverse of infectious duration) [$\text{time}^{-1}$] |
| $\mu_m$ | Per capita background mosquito mortality and birth rate (inverse of mosquito life expectancy) [$\text{time}^{-1}$] |
| $\mu$ | Per capita background host mortality and birth rate [$\text{time}^{-1}$] |
| $f$ | Mosquito fidelity (only bites imprinted host species if $f = 1$; bites host species $i$ with probability $\rho_i$ if $f = 0$) [unitless] |

**Table 3:** Estimates and range of parameter values used in model analysis and global sensitivity analysis.

| Parameter | Value | Range | Reference |
| --- | --- | --- | --- |
| $H$ | 1000 | (100, 1000) | Assumed |
| $\rho_A$ | — | (0.01, 0.5) | Mwandawiro et al. (1999, 2000) |
| $\beta$ | 1 | (0, 1) | Assumed |
| $\alpha$ | 1/3 day<sup>-1</sup> | (1/6, 1/2) | Birley and Rajagopalan (1981) |
| $\gamma$ | 1/5 day<sup>-1</sup> | (2, 7) | Ilkal et al. (1988) |
| $\mu_m$ | 1/30 day<sup>-1</sup> | (1/35, 1/15) | Assumed |
| $\mu$ | 1/365 day<sup>-1</sup> | (1/1095, 1/90) | Assumed |

**Figure 3:**
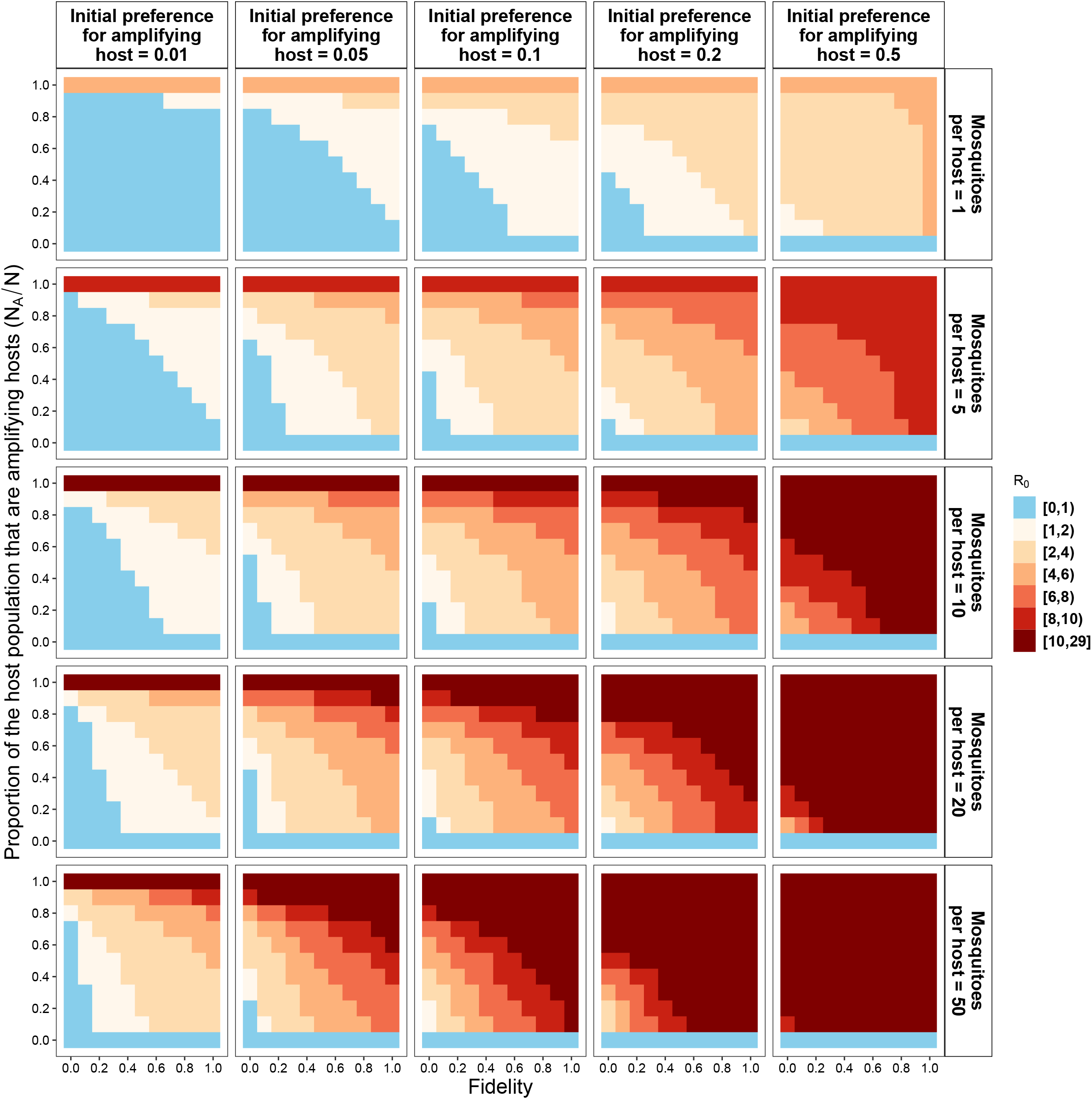
Basic reproduction number *R*_0_ values based on Equation S5 and values in Table 3. The top panel represents the initial preference for amplifying hosts, and the right panel is the mosquito-to-host ratio. Blue indicates regions where *R*_0_ *<* 1, i.e. there is no outbreak.

When there is a mixture of amplifying and dead-end hosts in the population, *R*_0_ increases with an increasing initial preference for amplifying hosts, fidelity, and mosquito-to-host ratio. When preference for the amplifying host species is 0.01, and given the fixed parameter values (Table 3), *R*_0_ is less than one when there are dead-end hosts in the population, except when the proportion of amplifying hosts *N*_*A*_*/N* exceeds 0.8 and the fidelity is sufficiently high (*f >* 0.6). With 50 mosquitoes per host and an amplifying host preference above 0.05, *R*_0_ is greater than 1, regardless of fidelity. However, an epidemic will not spread (*R*_0_ *<* 1) even if the mosquito-host ratio is as high as 50, provided that there are fewer amplifying hosts than dead-end hosts, the amplifying host preference less than 0.05 and *f* is less than 0.2. In the absence of fidelity and with a very low preference for the amplifying host, *R*_0_ is less than 1 if *N*_*A*_*/N* is less than 0.8, regardless of the number of mosquitoes per host. Therefore, fidelity can lead to *R*_0_ *>* 1 in circumstances where it would otherwise expected to be *<* 1.

### Transmission dynamics

Figure 4 shows the dynamics of infected host populations. In the absence of fidelity, an epidemic occurs only when there is an equal number of amplifying and dead-end hosts and the initial preference for the amplifying host is 0.2. In this scenario, the dead-end host population experiences a higher peak epidemic size than the amplifying host population, with the dead-end host reaching its peak first. Increasing fidelity results in larger peak epidemic sizes in both populations. However, beyond a certain level of fidelity, the peak epidemic size becomes larger in amplifying host populations than in dead-end populations. Perfect fidelity prevents epidemics in dead-end hosts, as those initially feeding on amplifying hosts maintain their preference.

**Figure 4:**
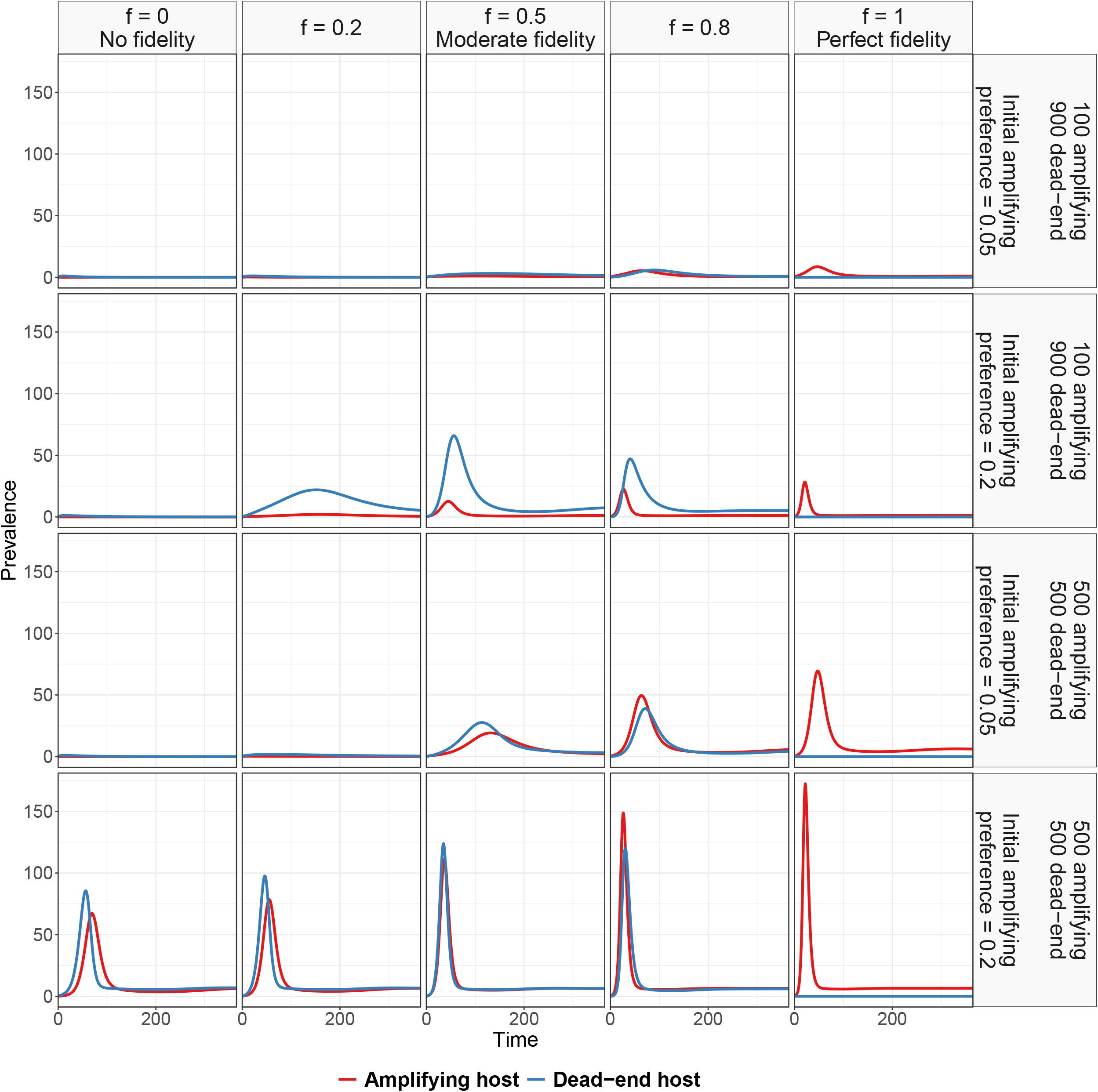
Dynamics of infected amplifying and dead-end host assuming 5 mosquitoes per host and fixed parameter values in Table 3. The right panel indicates host composition and initial preference for amplifying hosts, while the top panel indicates different fidelity values. While outbreak size in amplifying hosts increases with fidelity, less than perfect fidelity leads to larger outbreaks in the dead-end host.

In the amplifying host population, the epidemic size increases with increasing fidelity. However, in the dead-end population, the impact of fidelity varies based on initial preference and host composition. For example, when the amplifying to dead-end host ratio is 1, the peak epidemic size increases with increasing fidelity and initial preference. However, when there are 100 amplifying hosts and 900 dead-end hosts, and the initial preference is 0.2, the peak epidemic size increases with increasing fidelity values up to moderate fidelity, beyond which it decreases again.

### Comparison of relative *R*_0_ between different mosquito species

We used experiments conducted by Mwandawiro et al. (2000) to estimate the initial preference for amplifying host for the three mosquito species competent for JEV: *Cx. Tritaeniorhynchus* (*ρ*_*A*_ ≈ 0.05), *Cx. gelidus* (*ρ*_*A*_ ≈ 0.22), and *Cx. vishui* (*ρ*_*A*_ ≈ 0.15). Subsequently, we used the mosquito biting behaviour described in Figure 1 to estimate the fidelity values of each mosquito species using maximum likelihood estimation (see Figure S2). The fidelity estimates and their 95% confidence interval are 0.53 [0.31, 0.74] for *Cx. tritaeniorhynchus*, 0.57 [0.26, 0.88] for *Cx. gelidus*, and 0.75 [0.49, 1.00]) for *Cx. vishui*.

Each of the three JEV vectors has the potential to trigger an outbreak in a population consisting of only 1% amplifying host, with an average of 5 mosquitoes per host (Figure 5). Note that the value of *R*_0_ will depend on the extrinsic incubation period of mosquitoes, which is not considered here. In addition, other parameters used in this analysis are held constant under relatively lenient conditions (see caption in Figure 5). However, we performed a global sensitivity analysis to assess the variability of *R*_0_ across all parameter ranges, as shown in Figure S3.

**Figure 5:**
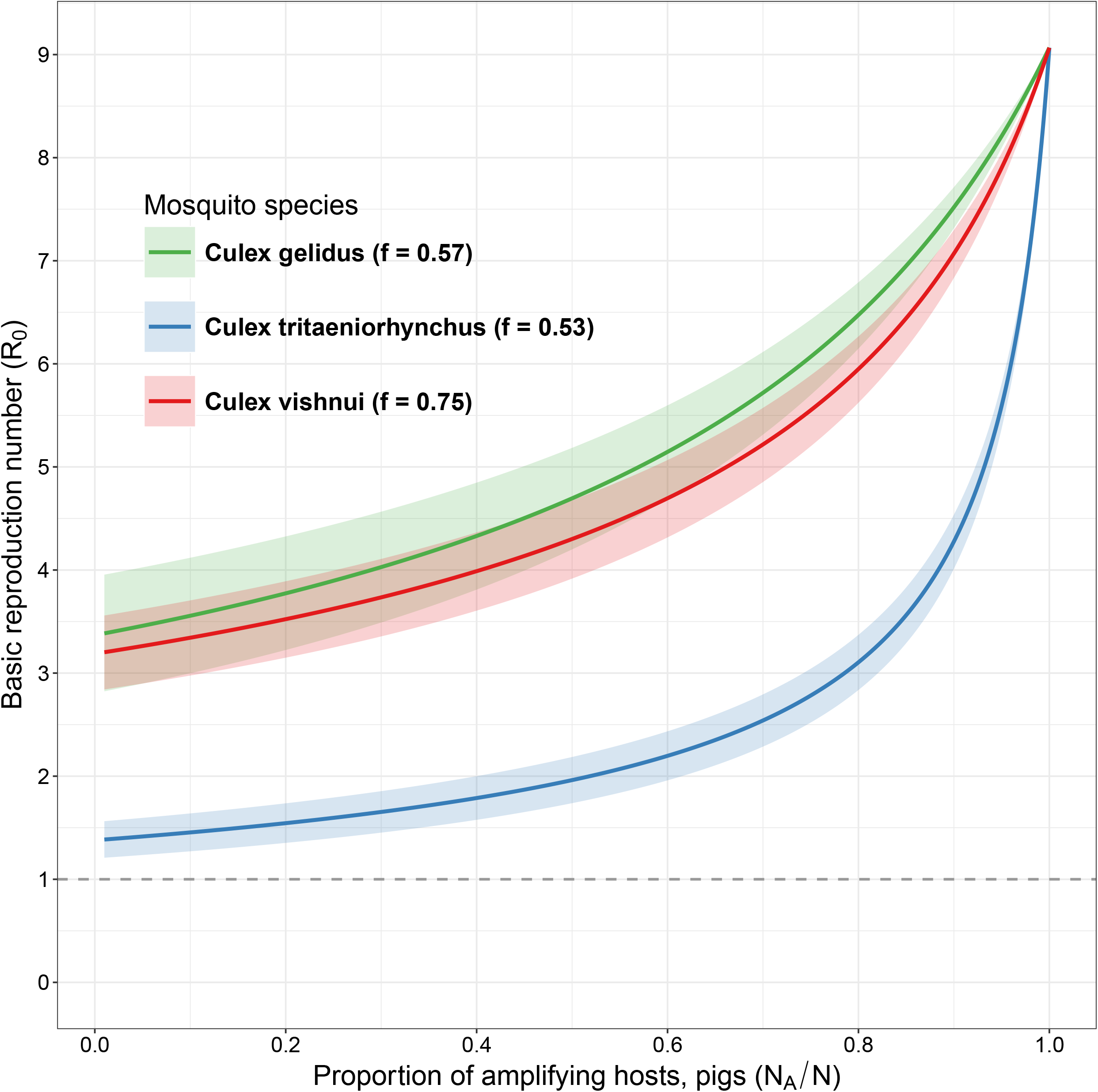
Basic reproduction number (*R*_0_) for *Cx. tritaeniorhynchus, Cx. gelidus*, and *Cx. vishui* based on varying host composition 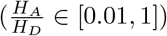. Initial preference for amplifying host *ρ*_*A*_ and fidelity values *f* estimated from Mwandawiro et al. (2000): 0.05 and 0.53 (0.31, 0.74) for *Cx. tritaeniorhynchus*, 0.22 and 0.57 (0.26, 0.88) for *Cx. gelidus*, and 0.15 and 0.75 (0.49, 1.00) for *Cx. vishui*. Other parameters held constant: transmission probability—1, biting rate—1*/*3 day^−1^, mosquito mortality and birth rate — 1*/*30 day^−1^, host recovery rate—1*/*5 day^−1^, host mortality, birth rate—1*/*365 day^−1^, and mosquitoes-to-host ratio — 5.

*Cx. gelidus* has an initial preference of 0.22 for amplifying host species, while *Cx. tritaeniorhynchus* has an initial preference of 0.05. *Cx. gelidus* has a higher initial preference for amplifying host species compared to *Cx. tritaeniorhynchus*. Despite similar fidelity estimates, the potential for an outbreak is substantially higher in *Cx. gelidus*. On the other hand, for *Cx. gelidus* and *Cx. vishnui* with similar initial preferences, the potential for an outbreak remains comparable, even though the fidelity estimate of *Cx. vishnui* is substantially higher than that of *Cx. gelidus*.

## DISCUSSION

Here we have shown how vector memory of, and subsequent fidelity to, the host species they take a first blood meal from, can permit invasion and onward transmission of a zoonotic mosquito-borne virus in contexts where this would not be possible otherwise. This may include circumstances where vector species prefer dead-end hosts and where dead-end hosts are the most common vertebrate species. Although vector fidelity does not affect the overall proportion of blood meals on each host species, we have shown how it can lead to non-homogeneous mixing between the host and vector community; when a mosquito has fidelity for host species, those mosquitoes who first feed on amplifying hosts are more likely to take subsequent feeds from amplifying hosts.

In our analyses, we have used JEV as an example of the impact of fidelity on transmission. Our simulations demonstrate that in regions where pigs are rare relative to cattle, which are dead-end hosts but preferred by common JEV vectors, JEV could be maintained if vectors display sufficient host species fidelity. While we used the example of the role of pigs and cattle in JEV transmission, the findings may also extend to poultry. As highlighted by Lord et al. (2015) poultry are common in many regions where JEV occurs and produce sufficient viremia to infect mosquitoes, but their role in transmission has not been fully quantified. The extent to which fidelity may contribute to the role of poultry in JEV transmission is unknown. It is likely that both presence and abundance of ornithophilic mosquitoes and fidelity in more generalist vectors will contribute to determining what role poultry play in JEV transmission in a given ecological context. These findings have implications for any risk assessment of JEV spread to new areas that involves consideration of host and vector species community composition; transmission could be sustained in communities dominated by dead-end hosts because of vector host species fidelity.

There is empirical evidence that the fidelity of *Cx. tritaeniorhynchus* may be lower than that of *Cx. vishnui* and its preference for pigs may be lower than both *Cx. vishnui* and *Cx. gelidus*. This has implications for the relative role of each vector species in JEV transmission in regions where cattle are the dominant vertebrate host; *Cx. tritaeniorhynchus* may not always be the most important vector. More generally, it may be important to consider fidelity as well as vector abundance, competence and host preference when incriminating mosquito species in pathogen transmission.

Vector fidelity may be important to consider for other vector-borne disease systems. For example, there are contexts where zoophilic vectors are important in malaria transmission (Garrett-Jones et al., 1980; Gatton et al., 2013; Waite et al., 2017) and the role of cattle in transmission and control of malaria has been assessed. The presence of cattle can either exacerbate (zoopotentiation) or reduce transmission (zooprophylaxis), depending on the location of cattle with respect to humans and on vector feeding behaviour (Donnelly et al., 2015). While in some circumstances the presence of cattle may increase the incidence of malaria, their role in control could be enhanced by the use of insecticides. Waite et al. (2017) combined empirical studies with modelling to show that increasing vector mortality during feeding on non-human hosts can contribute to reducing malaria transmission. Ruiz-Castillo et al. (2022) also highlight potential strategies including insecticide-treated cattle. However, we are not aware of any analyses of malaria control using livestock that factor in the potential role of vector fidelity. Non-homogeneous mixing caused by fidelity could undermine interventions aimed at livestock. However, modelling to further explore this would be necessary.

In our model, we considered the transmission of JEV within two-host communities, thereby neglecting the potential influence of other host species that could serve as alternative sources of blood meals. However, our model is adaptable and can be extended to include a broader range of alternative hosts. In addition, we ignored some ecological factors that could affect the potential of outbreaks, including the seasonal variation in feeding preference, which could directly impact on mosquito fidelity behaviour. While we have explored simulation scenarios for a broad range of host preference and fidelity, our analysis of species specific outbreak risk (*R*_0_) relies on fidelity and host preference estimated from a single experimental study conducted by Mwandawiro et al. (2000). The scarcity of studies replicating this analysis highlights the need for increased attention to fidelity behaviour in mosquitoes within the mosquito-borne disease research community.

Our work contributes to the wider vector-borne pathogen modelling literature, which in the last decade has expanded beyond the original Ross-MacDonald framework. While obtaining species-level fidelity estimates for different vector species would require semi-field experiments and may often not be possible, and there are also spatial heterogeneities that likely affect transmission, here our point is to highlight the potential role of fidelity and for this to be considered as a potential drive of transmission in regions where host community composition would, in the first instance, suggest that transmission is not possible.

## ACKNOWLEDGEMENTS

The authors gratefully acknowledge Juliet Pulliam, who originally conceptualised and developed the original version of the model with Mike Boots.

## AUTHOR CONTRIBUTIONS

Conceptualisation: TAS, MB Methodology: TAS, JSL, AJW, MB Investigation: TAS, JSL, AJW, MBB, MB Visualisation: TAS, JSL, AJW Writing: TAS, JSL AJW Review: TAS, JSL, AJW, MBB, MB Supervision: JSL, MBB.

## CODE AVALAILABILITY

The code used in this study is available in a private GitHub repository at https://github.com/tijanisulaimon/mosquitoFidelityModel. Access to the repository is available upon request from the corresponding authors.

## SUPPLEMENTARY MATERIAL

### Derivation of basic reproduction number

Mathematically, *R*_0_ is derived from the largest eigenvalue of the Next Generation Matrix (NGM) *FV* ^−1^, where:

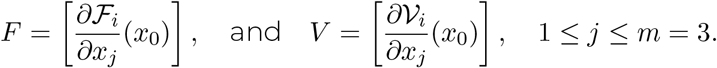

Here, *F*_*i*_(*x*) represents the rate of new infections appearing in infectious compartment *i* and 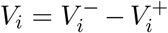 represents the net transition rate between compartment *i* and other infected compartments due to other means (van den Driessche and Watmough, 2002). Specifically, 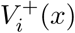 denotes the transfer rate into compartment *i*, while 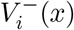 denotes the transfer rate out of compartment *i*.

From the simple model in 2, the disease-free equilibrium (DFE) is (*S*^*m*^, *I*^*m*^, *S*_*A*_, *I*_*A*_, *R*_*A*_, *S*_*D*_, *I*_*D*_, *R*_*D*_) = (*N*^*m*^, 0, *H*_*A*_, 0, 0, *H*_*D*_, 0, 0). The Next Generation Matrix (NGM) approach of van den Driessche and Watmough (2002) can be used to separate the infected equations into two matrices, ℱ and 𝒱 as follows:

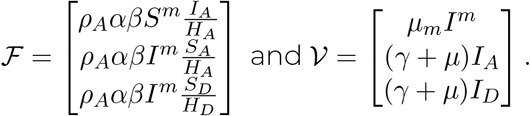

Lineariasing the matrices at DFE, we get:

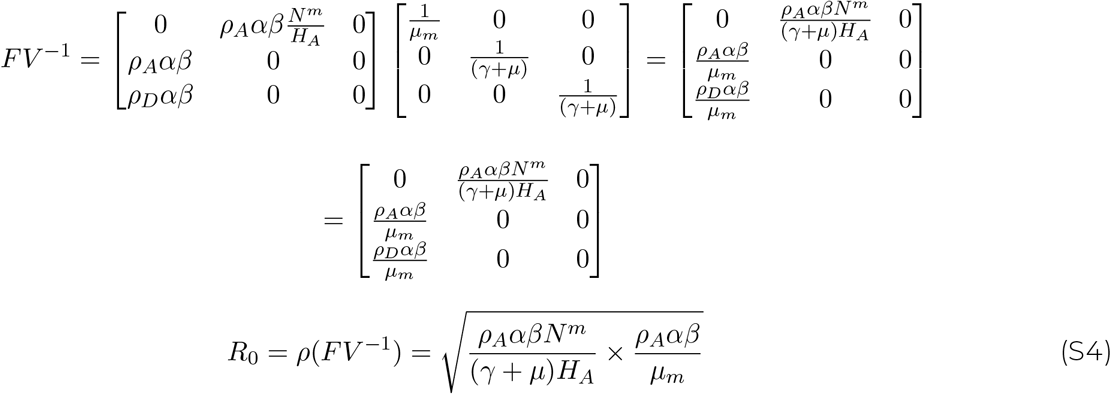

For the extended model in Equation 3, at DFE, 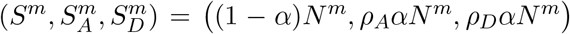, and the corresponding matrices ℱ and 𝒱 are:

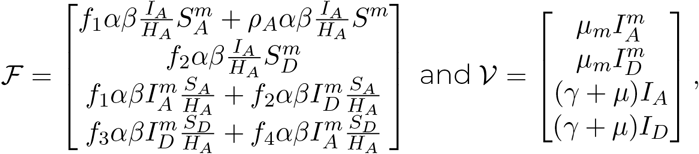

with *f*_1_ = [*f* + (1 − *f*)*ρ*_*A*_], *f*_2_ = [(1 − *f*)*ρ*_*A*_], *f*_3_ = [*f* + (1 − *f*)(1 − *ρ*_*A*_)], and *f*_4_ = [(1 − *f*)(1 − *ρ*_*A*_)].

Compute the linearised matrices at DFE,

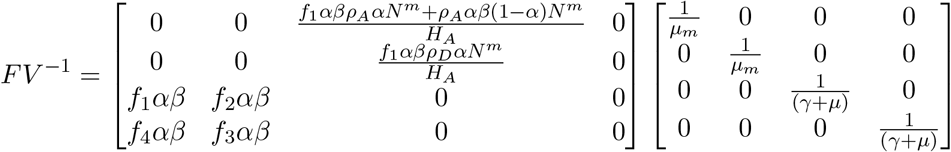

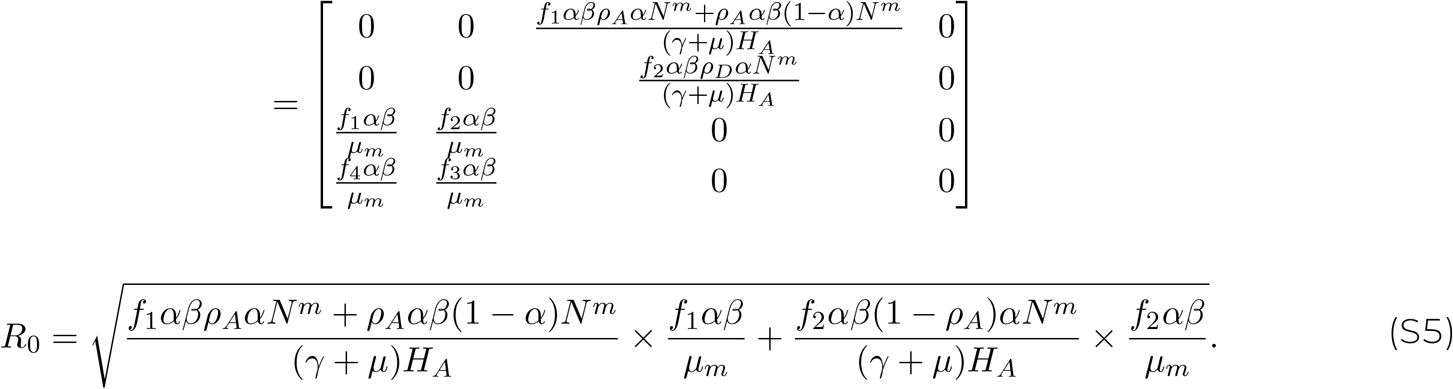

**Figure S1:**
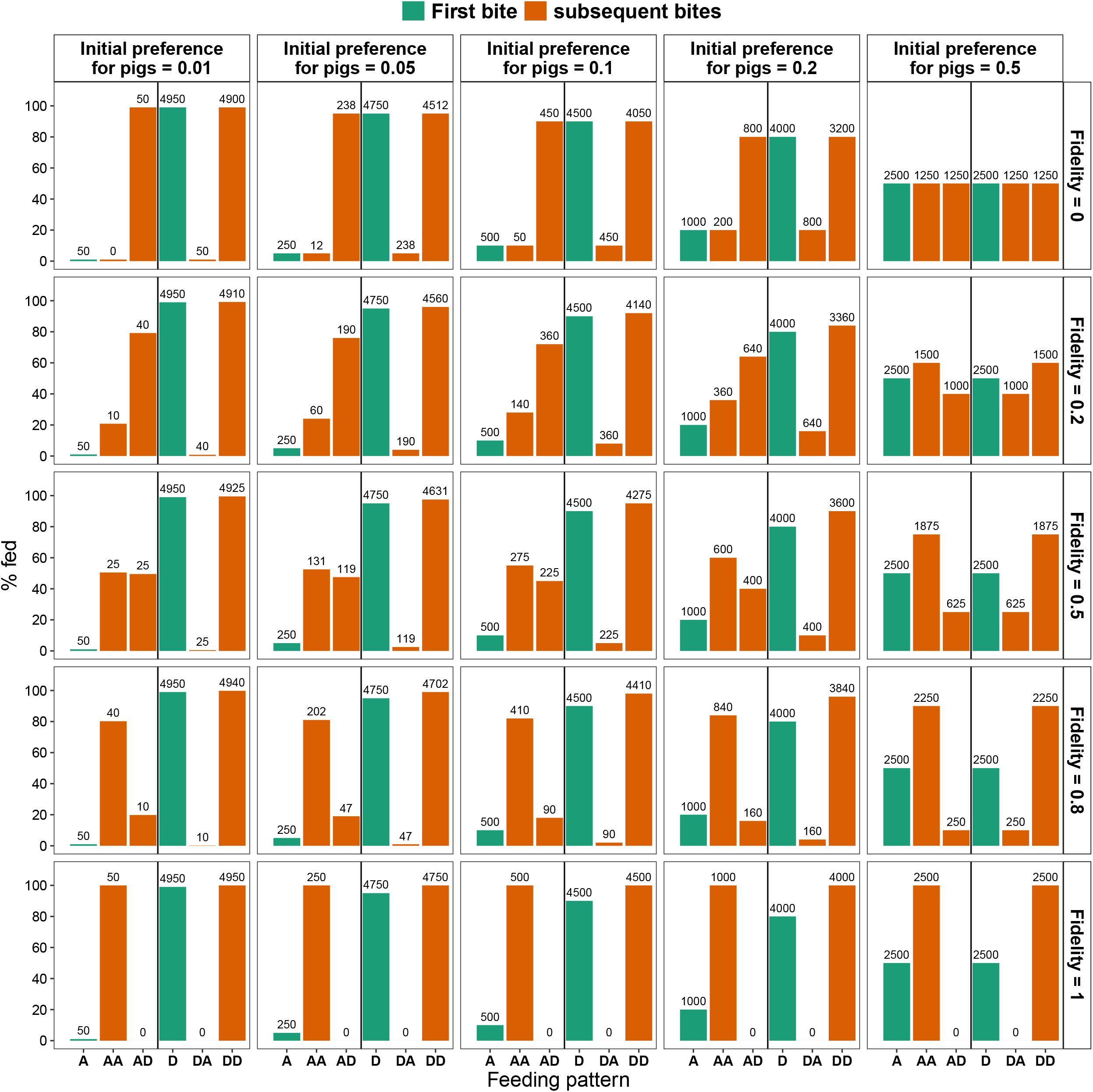
Variation of feeding patterns of mosquitoes with fidelity, in the instance where cows (C) and pigs (P) are available in equal proportions. The height of the bars indicates the proportion of mosquitoes that fed on host species (x-axis), with their corresponding numbers at the top. The fill indicates the proportion of first and subsequent bites. The top panel represents an increasing initial preference for pigs (left-to-right), while the right panel represents an increasing fidelity (top-to-bottom).

**Figure S2:**
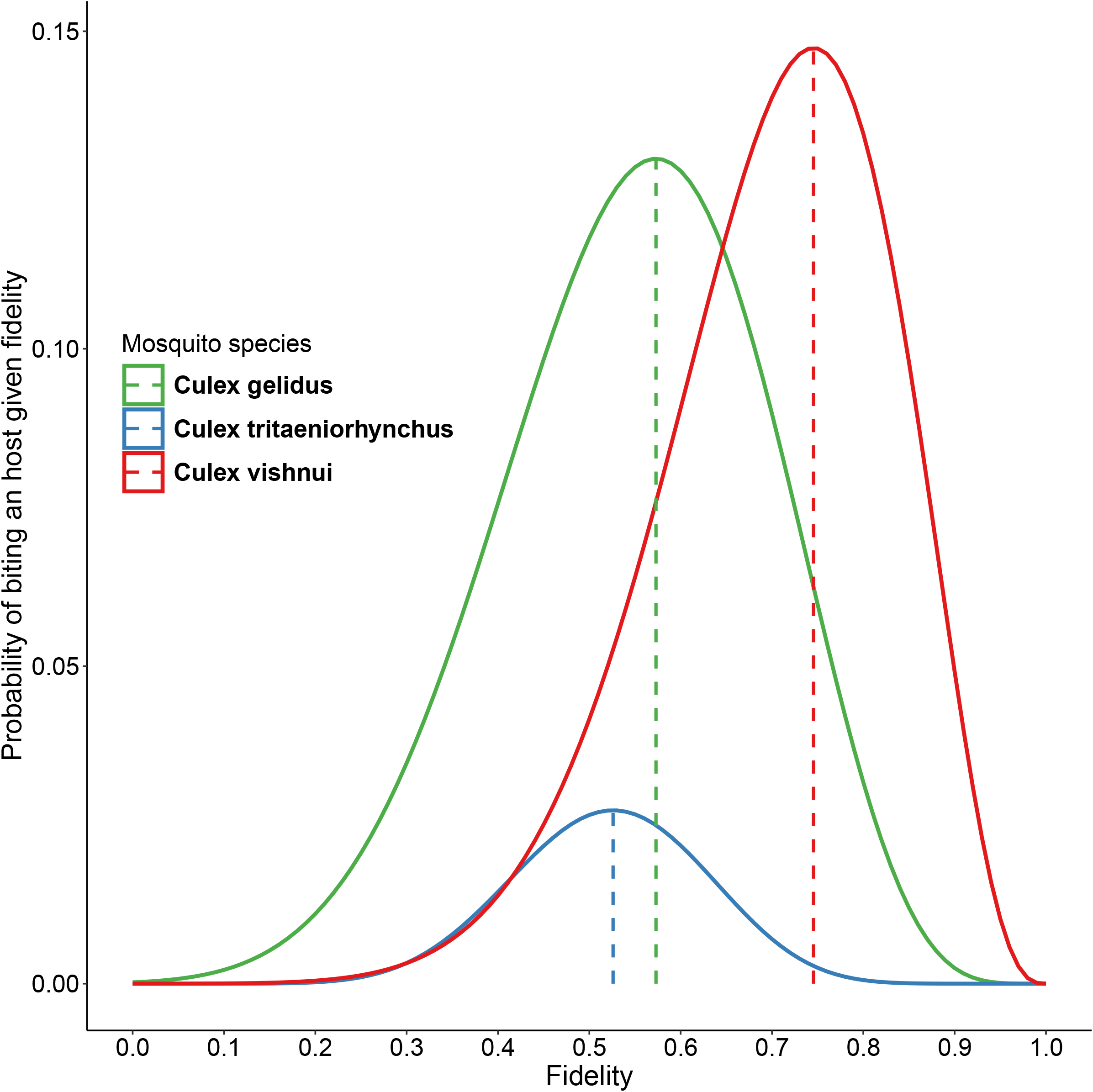
Fidelity values of three JE mosquitoes. Based on the experiment conducted by Mwandawiro et al. (2000), we estimated the initial preference for amplifying host for the three mosquito species competent for JEV: *Cx. tritaeniorhynchus* (*ρ*_*A*_ ≈ 0.05), *Cx. gelidus* (*ρ*_*A*_ ≈ 0.22), and *Cx. vishui* (*ρ*_*A*_ ≈ 0.15). Subsequently, we used the mosquito biting behaviour described in Figure 1 to estimate the fidelity values of each mosquito species using maximum likelihood estimation. This was performed in R using the “optim” function (R Core Team, 2024). The fidelity estimates obtained are *f* ≈ 0.53 (with a 95% confidence interval of [0.31, 0.74]) for *Cx. tritaeniorhynchus, f* ≈ 0.57 (95% confidence interval: [0.26, 0.88]) for *Cx. gelidus*, and *f* ≈ 0.75 (95% confidence interval: [0.49, 1.00]) for *Cx. vishui*.

**Figure S3:**
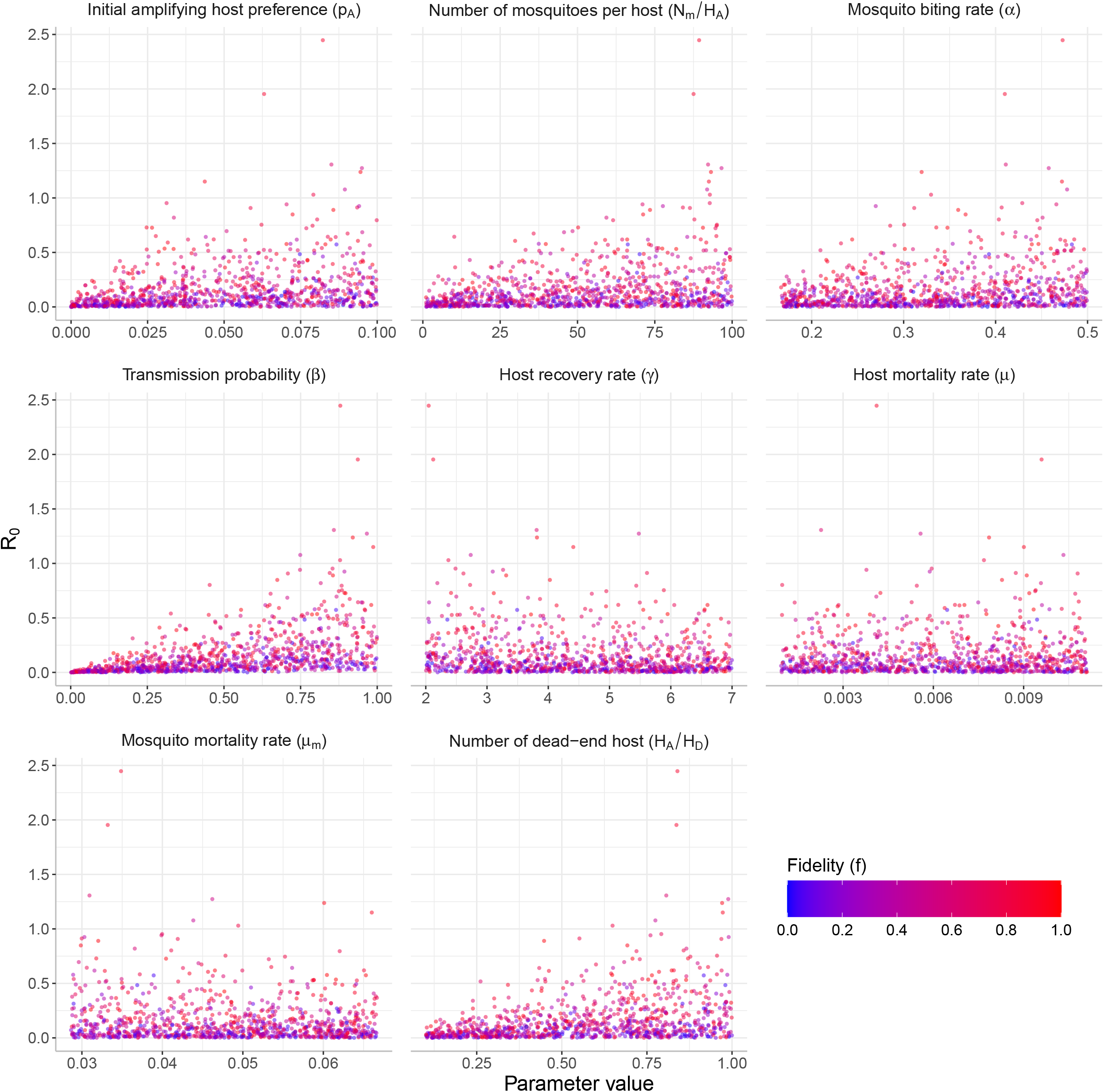
Global sensitivity analysis. To show the variability of *R*_0_ with respect to all parameter ranges, we used random Latin hypercube design to simulate 1 000 parameter sets with the ranges of parameters from Table 3, which were then used to calculate *R*_0_ values.

